# Probabilistic representation in human visual cortex reflects uncertainty in serial decisions

**DOI:** 10.1101/671958

**Authors:** R.S. van Bergen, J.F.M. Jehee

## Abstract

How does the brain represent the reliability of its sensory evidence? Here, we test whether sensory uncertainty is encoded in cortical population activity as the width of a probability distribution – a hypothesis that lies at the heart of Bayesian models of neural coding. We probe the neural representation of uncertainty by capitalizing on a well-known behavioral bias called serial dependence. Human observers of either sex reported the orientation of stimuli presented in sequence, while activity in visual cortex was measured with fMRI. We decoded probability distributions from population-level activity and found that serial dependence effects in behavior are consistent with a statistically advantageous sensory integration strategy, in which uncertain sensory information is given less weight. More fundamentally, our results suggest that probability distributions decoded from human visual cortex reflect the sensory uncertainty that observers rely on in their decisions, providing critical evidence for Bayesian theories of perception.

## Introduction

Many of our day-to-day decisions are plagued by uncertainty. For example, when crossing a road, it is near impossible to estimate the trajectories of oncoming traffic with absolute precision. Rather, our perceptual estimates of vehicle trajectory patterns are *uncertain*, due to variability and ambiguity in the sensory inputs. Numerous behavioral studies have shown that human observers utilize knowledge of sensory uncertainty when making perceptual decisions (Jacobs and Fine, 1999; Ernst and Banks, 2002; Battaglia et al., 2003; Knill and Saunders, 2003; Alais and Burr, 2004), but how does the brain represent this uncertainty in our sensory estimates?

Bayesian theories of neural coding propose that uncertainty is represented in neural activity as the width of a probability distribution (Zemel et al., 1998; Anastasio et al., 2000; Hoyer and Hyvärinen, 2003; Jazayeri and Movshon, 2006; Ma et al., 2006; Fiser et al., 2010). That is, neural population activity is typically consistent with a whole range of stimuli, rather than a single-valued estimate. Mathematically, this range can be represented as a probability distribution over stimulus values. The width of the distribution can be taken as a measure of the degree of uncertainty contained in the population response – and it is this probabilistic information, these theories propose, that observers utilize in their decision-making. Using functional MRI and a probabilistic decoding analysis, we have previously shown that probability distributions reflecting sensory uncertainty can reliably be extracted from the human visual cortex (van Bergen et al., 2015; van Bergen and Jehee, 2018). However, it remains to be determined whether observers also utilize this representation of uncertainty when making decisions. Although indirect evidence appears consistent with this notion (van Bergen et al., 2015), unequivocal support for the hypothesis is still lacking.

To study the neural code for uncertainty, we capitalize on a well-studied behavioral bias called serial dependence (Cicchini et al., 2014; Fischer and Whitney, 2014; Liberman et al., 2014): when a stimulus is embedded in a sequence, human observers tend to judge it as being more similar to previously seen stimuli than it really is (but see Gibson and Radner, 1937; Chopin and Mamassian, 2012; Fritsche et al., 2017). Serial dependence is particularly well suited to the issue at hand because it appears to reflect a statistically advantageous sensory integration strategy (Cicchini et al., 2014; Fischer and Whitney, 2014). That is, considering that the natural environment is largely stable across time (Dong and Atick, 1995), the statistically ideal observer integrates past and present sensory inputs, weighting each by its associated uncertainty. This integration process not only results in more accurate behavior, but also biases behavioral estimates towards previously seen stimuli. Critically, for the ideal observer, the magnitude of the biasing effect depends on the degree of sensory uncertainty. This, then, yields a straightforward prediction that we leverage in the present study: if the decoded probability distributions reflect the uncertainty that is utilized in decisions (as hypothesized by probabilistic theories of neural coding), then decoded uncertainty should be directly linked to the magnitude of the serial dependence bias observed in human behavior.

Our goals for the current study are therefore twofold: to investigate the computational goals underlying serial dependence in perception, and in doing so, probe the neural representation of sensory uncertainty. We first describe a theoretically ideal observer who infers the stimulus from noisy sensory inputs in a temporally predictable environment. We then test the model’s predictions against behavioral and brain data obtained from human observers, using a probabilistic decoding analysis to characterize the degree of uncertainty in cortical stimulus representations (areas V1-V3). We find that serial dependence biases in human behavior are consistent with a statistically ideal inference process that not only takes into account the temporal stability of the natural environment, but also respects the degree of uncertainty associated with each successive input. More fundamentally, these results suggest that probability distributions decoded from population activity in human cortex reflect the uncertainty that observers rely on when making perceptual decisions, providing critical evidence for Bayesian theories of perception.

## Materials and Methods

### Participants

Eighteen healthy, adult volunteers (seven female, eleven male, aged 22-31 years) participated in this study, which was approved by the Radboud University Institutional Review Board. Subjects provided written and informed consent before participation.

### Data acquisition

We only briefly describe fMRI data acquisition and preprocessing procedures here. For a full description, please see (van Bergen et al., 2015), in which data were previously analyzed for a different purpose. The MRI data were acquired at the Donders Center for Cognitive Neuroimaging, using a Siemens 3T Magnetom scanner equipped with an eight-channel occipital receiver coil. Each scan session started with the collection of a high-resolution T1-weighted anatomical scan, using a magnetization-prepared rapid gradient-echo protocol (MPRAGE, 1-mm isotropic voxels, FOV 256 × 256). Functional images covered all of occipital and some of posterior parietal and temporal cortex, in 30 slices oriented perpendicular to the calcarine sulcus, and were scanned with a T2*-weighted gradient-echo echoplanar imaging sequence (TR 2000 ms, TE 30 ms, FOV 64 × 64, 2.2 mm isotropic voxels, flip angle 90°).

### Experimental design & stimuli

Visual activity and behavioral responses were measured while participants viewed orientation stimuli inside an fMRI scanner. Throughout each run, participants were instructed to maintain visual fixation at a central black-and-white bull’s eye target (radius: 0.25 deg.). A run consisted of 18 orientation trials presented in sequence, with a 4-s fixation period at the start and end of the run. Each orientation trial was separated by a 4-s inter-trial interval. A trial (**Fig. 1**) started with the presentation of a sinusoidal grating (duration: 1.5 s, spatial frequency: 1 cycle/deg., randomized spatial phase, 2 Hz sinusoidal contrast modulation, peak contrast: 10%) presented inside an annulus surrounding fixation (inner radius 1.5 deg., outer radius: 7.5 deg., grating contrast decreased linearly to 0 over the outer and inner 0.5 deg. radius of the annulus). Grating orientation was determined (pseudo-)randomly to ensure an approximately even sampling of orientation space (a prerequisite for an unbiased decoding analysis). After a brief interval (6.5 s), a black bar (width: 0.1 deg., length: 2.8 deg.) appeared at the center of the screen at an initially random orientation. Participants reported the orientation of the stimulus by rotating the bar, pressing separate buttons for clockwise or counter-clockwise rotation on an MRI-compatible button box. The bar remained on screen for a total of 4 s, and started to fade-in to the background during the final 1 s of this window to indicate the approaching response deadline.

**Figure 1:**
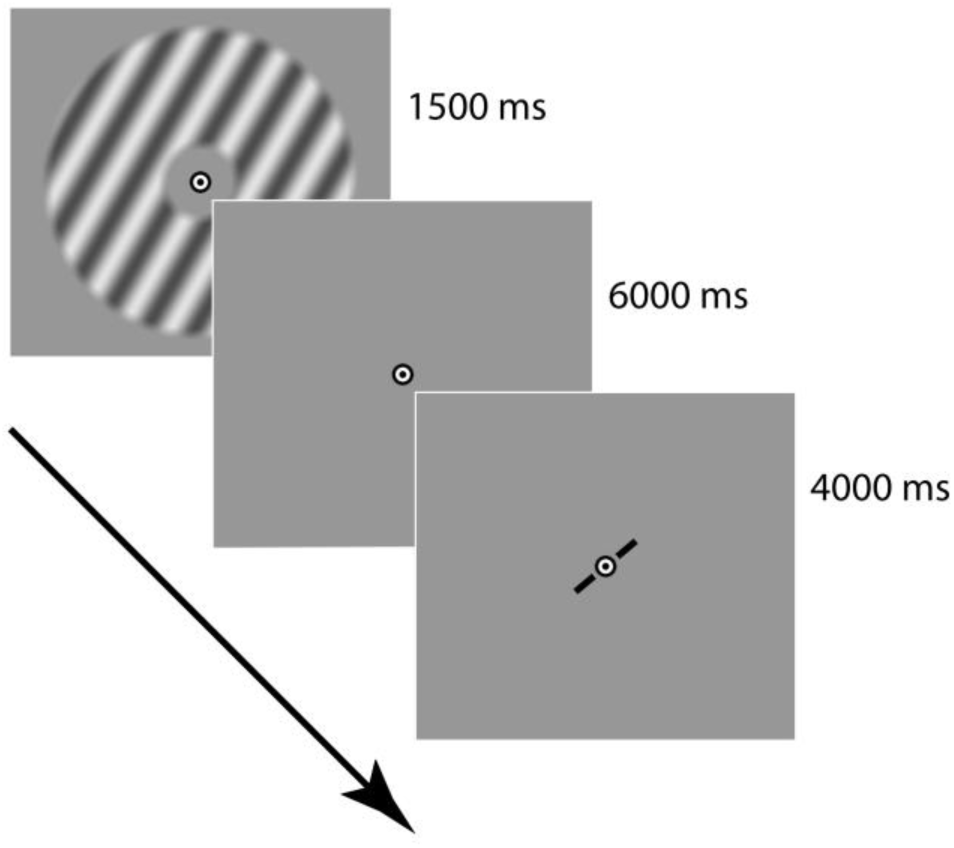
Trial structure. Each trial in the experiment started with the presentation of a stimulus, followed by a fixation interval, and then a response window. Trials were separated by a 4000 ms inter-trial interval. Participants adjusted the orientation of a centrally presented bar to match the previously seen stimulus orientation. The stimulus and response bar are not drawn to true scale and contrast.

Participants completed 10-18 runs of the orientation task, totaling 180-324 trials, as well as two visual localizer runs within the same scan session. In the localizer runs, flickering checkerboard patterns were presented in 12-s blocks, within the same annular aperture as the orientation stimuli (contrast: 100%, check size: 0.5 deg., 10 Hz updating of the random checkerboard pattern), and interleaved with fixation blocks of equal duration. In a separate scan session, retinotopic maps of visual cortex were acquired using standard retinotopic mapping procedures (Sereno et al., 1995; DeYoe et al., 1996; Engel et al., 1997).

Visual stimuli in the fMRI scanner were displayed on a rear-projection screen by a luminance-calibrated EIKI projector (resolution 1024 × 768 pixels, 60 Hz refresh rate), and viewed by participants through a mirror mounted on the head coil. Stimuli were generated by a Macbook Pro computer running MATLAB and the Psychophysics Toolbox (Brainard, 1997; Pelli, 1997).

### fMRI data pre-processing and regions of interest

Functional images were motion-corrected using FSL’s MCFLIRT (Jenkinson et al., 2002), and filtered in the temporal domain to remove slow drifts in the BOLD signal (high-pass cut-off: 40 s). Residual motion-induced fluctuations in BOLD signal were removed through linear regression (18 motion regressors, constructed from the motion estimates produced by MCFLIRT). Functional images were registered to a previously collected anatomical reference scan using FreeSurfer (Fischl et al., 1999). All functional data were analyzed in native space.

Regions of interest (ROIs; V1, V2 and V3) were defined on the reconstructed cortical surface using standard procedures (Sereno et al., 1995; DeYoe et al., 1996; Engel et al., 1997). Within each ROI, all voxels were selected that were activated by the functional localizer stimulus at a lenient statistical threshold (*p* < 0.05, uncorrected). Control analyses verified that our results were highly robust to changes in the number of voxels selected for analysis (i.e. a range of voxel selection thresholds between p < 0.005 and p < 0.2).

Each voxel’s time series was z-normalized using the corresponding time points of all trials in a given run. Activation patterns for each trial were defined by first adding a 4-s temporal shift (to account for hemodynamic delay), and then averaging together the first 4 s of each trial. This selected time window was relatively short (4 s) so as to make sure that activity from the behavioral response window was excluded from analysis.

### Decoding analysis

A generative model-based, probabilistic decoding algorithm was used to characterize the trial-by-trial uncertainty in cortical stimulus representations (van Bergen et al., 2015; van Bergen and Jehee, 2018). The analysis assumes that voxel activations follow a multivariate Normal distribution around a stimulus-dependent mean, where the latter is described by the orientation tuning function of each voxel. Voxel tuning curves were modeled as a linear combination **W*f***(*s*) of eight bell-shaped basis functions (Brouwer and Heeger, 2011):

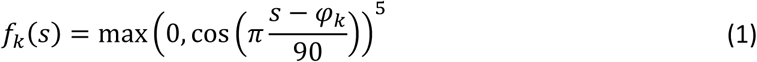

where *s* is the orientation of the stimulus and *φ*_*k*_ is the preferred orientation of the *k*-th basis function (both in °). Basis functions are weighed by coefficients **W** = {*W*_*ik*_} for each voxel *i* and basis function *k*. The covariance of the noise distribution around the voxel tuning functions is modeled as:

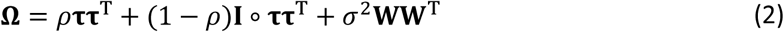

This noise covariance matrix is a combination of three components. The first component is a diagonal of independent noise variances **τ** = {*τ*_*i*_}, where the variance of voxel *i* is given by 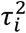. The second component is a constant that models fluctuations in signal shared by all voxels, and is described by a parameter *ρ*. Finally, the term *σ*^2^**WW**^T^ models noise, with variance *σ*^2^, that is shared between voxels with similar orientation tuning preferences.

Together, the orientation tuning curves and the noise covariance structure make up the decoder’s generative model. This model specifies the generative distribution *p*(**b**|*s*): the probability that a certain stimulus *s* will evoke an activation pattern **b**. Thus, this distribution is given by a multivariate Normal with mean **W*f***(*s*) and covariance **Ω**:

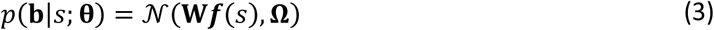

where **θ** = {**W, τ**, *ρ, σ*} are the parameters of the generative model.

Model parameters were estimated using the fMRI activation patterns in a leave-one-run-out cross-validation procedure. Data were divided into a training dataset (consisting of data from all but one fMRI run) and a testing dataset (consisting of data from the remaining run). The parameters of the generative model were fit to the training data using a two-step estimation procedure. In the first step of this procedure, tuning weights **W** were estimated by ordinary least-squares regression. In the second step of the parameter-estimation procedure, noise covariance parameters (**τ**, *ρ* and *σ*) were estimated by numerically maximizing their likelihood. See (van Bergen et al., 2015) for further information regarding these fitting procedures.

After fitting the model to the training data set. We tested the model on the held-out (independent) testing data set. For each test trial, we calculated the posterior distribution over stimulus orientation, conditioned on the estimated model parameters 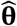. Following Bayes’ rule, the posterior distribution is given by:

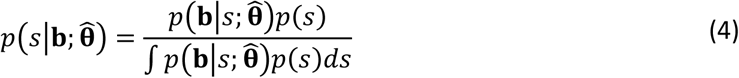

The stimulus prior *p*(*s*) was flat (reflecting the uniform distribution of orientation stimuli), and the normalizing constant in the denominator was calculated numerically. The (circular) mean of the posterior function served as an estimate of the presented orientation on that trial, while the (circular) standard deviation was taken as a measure of the degree of uncertainty in the orientation estimate. The leave-one-run-out cross-validation procedure was repeated until each run of fMRI data had served as the test set once. To ensure that decoded uncertainty reflected trial-by-trial, rather than orientation-related (Appelle, 1972; Furmanski and Engel, 2000; van Bergen et al., 2015), fluctuations in the precision of a cortical representation, orientation-dependent effects on uncertainty were removed via linear regression (see also (van Bergen et al., 2015)).

### Behavioral data

The observer’s behavioral error on a given trial was computed as the acute-angle difference between the reported and presented orientation, with positive angles corresponding to clockwise deviations. Participants generally performed well on the task, with errors in their orientation reports averaging 6.29 +/-0.25 (mean +/-SEM across observers).

To analyze serial dependence effects, data were first corrected for a repulsion bias away from the cardinal axes (van Bergen et al., 2015) by fitting each observer’s behavioral errors with a Gaussian-Uniform mixture distribution (Zhang and Luck, 2008) around a 4-degree polynomial function of stimulus orientation. The residuals from this polynomial fit were used in the remaining analyses. Trials for which the probability of the fitted uniform distribution was larger than p = 0.5 were assumed to be random guesses by the participant, and excluded from further analysis (0-3 trials out of 180-324 per observer).

The serial dependence effect in participant behavior was characterized by means of a group-average bias curve, which was obtained as follows. For each individual observer, the acute-angle difference between current and previous stimulus orientation was first calculated for all consecutive trials in each run. The mean behavioral error across trials was then computed for each orientation difference in the experiment. The data were smoothed with a moving average filter (window width = 20°) to create observer-specific serial dependence curves across orientation differences. Finally, the curves were averaged across observers. Procedures were identical for a control analysis in which serial dependence biases were computed with respect to the behaviorally reported, rather than presented, orientation on the previous trial.

Many of our analyses leveraged the symmetry of the serial dependence effect by collapsing the serial dependence curve onto one side of the graph. This was achieved by mirroring each participant’s curve in the origin, and then averaging the mirrored and original curves together, before computing the group-average.

### Ideal observer model

Theoretical predictions were quantified using an ideal observer approach. Specifically, the observer model started with the assumption that the observer takes a sensory measurement from the environment at time *t*. The measurement is noisy: there is no one-to-one mapping between the external stimulus (*s*^(*t*)^) and internal measurement (*m*^(*t*)^). Rather, the relationship between stimulus and measurement is described by a probability distribution *p*(*m*^(*t*)^|*s*^(*t*)^). The observer’s task is to infer which stimulus (orientation) is presented. The ideal observer uses the *generative model*, i.e. *p*(*m*^(*t*)^|*s*^(*t*)^), to determine which stimulus most likely caused the sensory measurement. Specifically, the observer inverts the generative model, reasoning backwards from the internal measurement to likely causes, using Bayes’ rule:

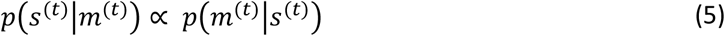

The resulting posterior distribution describes for every possible stimulus orientation, the probability that it caused the internal measurement. The peak of the distribution defines the most likely stimulus, while the distribution’s width (or variance) can be taken as a measure of the degree of uncertainty in this orientation estimate. Representing knowledge probabilistically is advantageous because it enables the observer to express the reliability of their sensory measurements, and utilize this information in further computations.

We continue with the notion that the natural environment is fairly stable over time (Dong and Atick, 1995). Specifically, we analyzed the temporal orientation statistics in a large range of natural videos, and found that natural changes in orientation over small periods of time are well approximated by a mixture distribution of a central peak and a uniform baseline (**Fig. 2**):

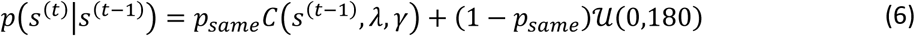

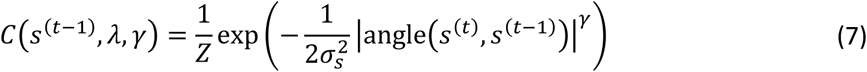

where *Z* is a normalization constant that was computed numerically. This transition distribution describes the probability that a stimulus at time *t* has orientation *s*^(*t*)^, given that the previous stimulus had orientation *s*^(*t*–1)^. The distribution consists of two components. The first component, a central peak, models stability in the environment, and is centered on no change. The width and kurtosis of this central peak are controlled by parameters *σ*_*s*_ and *γ*, respectively. When *γ* = 2, the distribution is Gaussian and the width parameter *σ*_*s*_ corresponds to the standard deviation of the central peak. For other values of *γ*, the width parameter is not easily interpretable, which is why we report numerically computed values of the distribution’s standard deviation and kurtosis in most of our analyses.

**Figure 2:**
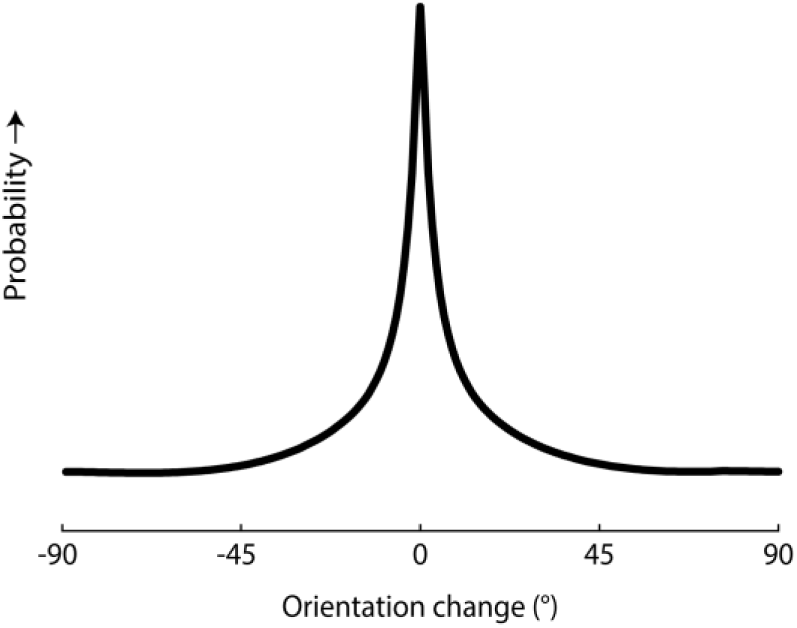
Distribution of orientation change over time in natural videos. Two databases of natural videos (Kayser et al., 2003; Betsch et al., 2004; Dorr et al., 2010) were analyzed for their orientation content as it evolved over time. Each video database was analyzed at three different spatial scales (lower, middle and upper third of the spatial frequency spectrum) and for three different temporal intervals (200, 1000 and 10,000 ms). Within each spatial scale, temporal interval and database, the change in orientation content across frames was computed for each location in the image, and orientation changes were tabulated across image locations and frames. See **Extended data figure 2-1**, for more details regarding the analysis procedures and parameter values, as well as separate figures for spatial scales, temporal intervals and video databases. Shown here is the average distribution of orientation changes across scales, intervals and databases. The distribution combines a flat baseline with a central peak, which was approximated by a mixture distribution of a central peak and a uniform baseline in the naturalistic observer model.

The second component in the mixture, a uniform distribution, accounts for the fact that sometimes (with probability 1 – *p*_*same*_) observers do encounter sudden and unpredictable changes in the environment (e.g., a previously seen bird has flown away). This component describes prior knowledge about the probability of orientation stimuli, regardless of stimulus history. For simplicity, we assume the prior to be uniform, but it could easily be replaced by, for instance, a distribution favoring cardinal orientations (Girshick et al., 2011) without affecting our general conclusions.

In a stable environment (as we assume here), it is advantageous for the observer to form predictions about the stimulus based on previous sensory measurements. Combining predictions with information from the current measurement tends to produce more accurate stimulus estimates than those based on current sensory information alone. To compute a prediction, 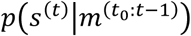, the ideal observer uses knowledge about the statistical structure of the environment, and combines this statistical knowledge with information obtained from previous observations:

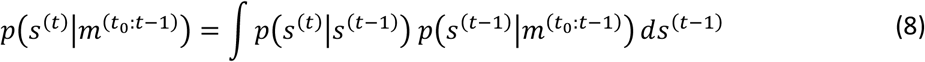

Thus, the prediction is obtained by convolving the distribution of knowledge about the previous stimulus 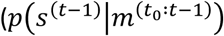, with the distribution of changes that can occur between two consecutive measurements (*p*(*s*^(*t*)^|*s*^(*t*–1)^)). Note that, due to the recursive nature of the ideal decision process, the distribution over previous stimuli (indirectly) reflects all of the information acquired since time *t*_0_, the starting point of the inference process.

The ideal observer combines this prediction with knowledge deduced from the current sensory measurement, resulting in a new distribution:

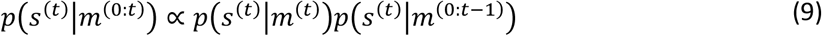

This final distribution represents the observer’s belief, expressed as probabilities, about the orientation of the currently viewed stimulus. This belief is based on all the information available to the observer at time *t*, including information obtained from both current and previous sensory observations. We assume that the observer reports the mean of this distribution as their best estimate of the viewed stimulus orientation. In the following, we refer to this observer model as the ‘naturalistic’ observer.

### Simulations

Behavioral orientation estimates of the naturalistic observer were obtained using a sequence of 10,000 orientation stimuli with naturalistic temporal statistics. Specifically, for each simulated trial *t*, a stimulus orientation *s*^(*t*)^ was randomly drawn from equation (6), with *p*_*same*_ = 0.9, *σ*_*s*_ = 10 and *γ* = 2, except for the initial stimulus at *t* = *t*_0_, which was drawn from a uniform distribution between [0, 180]°. Each stimulus orientation *s*^(*t*)^ evokes a noisy sensory measurement. Specifically, the sensory measurement on trial *t* was drawn at random from a Circular Gaussian (the sensory measurement distribution *p*(*m*^(*t*)^|*s*^(*t*)^)) centered on the true stimulus orientation *s*^(*t*)^:

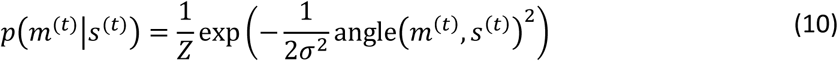

where *Z* is a normalization constant that was computed numerically, and *σ*^2^ is the variance of the sensory noise. Thus, the sensory measurement for a given orientation *s*^(*t*)^ fluctuates across trials due to noise. The noise parameter of the distribution varied between weak and strong levels of noise: on a random 50% of trials, the measurement distribution had a standard deviation of *σ* = 5° (weak sensory noise), while on the remaining trials this was 10° (strong sensory noise). Probabilistic inference of the most likely stimulus orientation on each trial proceeded with full knowledge of parameters values and according to the algorithm laid out in equations (5-9), except that on the first trial (*t* = *t*_0_), there was no prediction available from a preceding trial.

We compared naturalistic observer performance to the simulated behavioral orientation estimates of three alternative observers. Each of these observers lacked a component of the ideal inference process. The first observer ignored past sensory responses altogether. This observer was simulated by skipping the prediction step in the ideal observer algorithm, such that the observer’s perceptual knowledge reflected only the sensory input on the current trial. The second observer ignored trial-by-trial fluctuations in sensory uncertainty. This observer (incorrectly) assumed that the sensory posterior *p*(*s*^(*t*)^|*m*^(*t*)^) had a constant width of 7.9° on each trial (the optimal uncertainty in this scenario for an observer assuming constant uncertainty). The third alternative observer used incorrect assumptions about the temporal orientation statistics of the environment. While temporal correlations were determined by *p*_*same*_ = 0.9 in the simulated natural environment, this observer instead assumed a value of *p*_*same*_ = 1. Effectively, this model observer ignored the difference in orientation between consecutive observations, and simply averaged the two while taking into account their uncertainty. We refer to these three non-ideal observers as the ‘naïve’, ‘uncertainty-blind’ and ‘temporally misinformed’ observers, respectively.

To examine the simulated observer’s behavior under experimental (i.e. non-naturalistic) conditions, we repeated the simulation procedures described above, but with stimuli randomly drawn from a uniform distribution independent across time (this is equivalent to setting *p*_*same*_ = 0 in equation (6)).

### Model fits

To ascertain the extent to which the computational models captured human behavior, parameter values were fitted to the serial dependence biases in participant behavior across the full range of orientation angles. For the first analysis, the models were fit to the participants’ group-average serial dependence curves across trials, assuming a constant level of sensory uncertainty (posterior width = *u*_*constant*_) across trials. This was a free parameter in the model, in addition to three parameters used to model temporal predictions, *p*_*same*_, *σ*_*s*_ and *γ*. To ensure plausible parameters values given their interpretation, these four parameters were constrained to 1 ≤ *u*_*constant*_ ≤ 60, 0 ≤ *p*_*same*_ ≤ 1, *σ*_*s*_ > 0 and *γ* > 0.8. For the temporally misinformed observer, parameters were fixed to *p*_*same*_ = 1 and *γ* = 2. The fitting procedure entailed finding the values that optimally explained human behavior within these ranges. For reasons of clarity, none of the observer models included parameters for additional sources of noise (e.g. motor noise), as including these would have no effect on the magnitude or shape of the serial dependence curve.

A crucial prediction of the naturalistic observer model is that the strength of the serial dependence effect should depend on the difference in sensory uncertainty between the current and previous trial. In a second analysis, we therefore divided each participant’s trials into two bins. The first bin contained trials for which uncertainty (decoded from cortical activity) was higher on the current than previous trial (*high* → *low uncertainty*), while the second bin consisted of trials for which uncertainty was higher on the previous than current trial (*low* → *high uncertainty*). We tested the degree to which uncertainty played a role in participant decisions by comparing how well their behavior was captured by two different observer models. The first observer model assumed a fixed level of uncertainty across trials (see also above), thus ignoring trial-to-trial fluctuations in the reliability of their sensory evidence. The second simulated observer accounted for across-trial variability in uncertainty when making decisions, by weighting each piece of evidence by its reliability. This model was fit to the data by assuming two bins or levels of uncertainty (*u*_*low*_ and *u*_*high*_), both bounded between [1, 60]°, and further constrained such that *u*_*low*_ < *u*_*high*_.

To fit these models, the sum of squared residuals between predicted and observed serial dependence curves was minimized numerically, using MATLAB’s `lsqcurvefit` algorithm. The goodness-of-fit of the model and parameter values was evaluated by computing the coefficient of determination (*R*^2^). The circular standard deviation and kurtosis of the central peak of the fitted transition distributions were computed numerically to facilitate direct comparison between the fitted distributions and those estimated from natural videos (**Fig. 2**).

### Statistical analysis

To determine whether the orientation reports of the human participants displayed a statistically significant serial dependence bias, and to appropriately account for the dependencies between data points introduced by smoothing, we used a cluster-based permutation test modeled after (Maris and Oostenveld, 2007). First, we defined the largest cluster of orientation differences with a positive serial dependence bias in the group-average of the behavioral data. This cluster was found to lie between orientation differences of [1-57]°. We then computed *t*-statistics for the bias at each orientation difference in the cluster. This was computed as the mean behavioral error across participants for that orientation difference, divided by the corresponding standard error. The size of the cluster was then quantified using the cumulative *t*-statistic, calculated as the sum of *t*-statistics across the cluster’s data points. To assess statistical significance, we compared its size to the cluster sizes in a simulated null distribution of data sets. This null distribution was generated by randomly permuting the observed data 10,000 times. Each permutation was computed by inverting the sign of a random portion of behavioral errors for each participant (a standard approach for a one-sample permutation test against the null hypothesis that the data have zero mean). The permuted data were smoothed and averaged across participants, following the same procedures as applied to the observed data. The largest cluster with a positive serial dependence bias was then determined in this permuted group-average, and its size (cumulative *t*-statistic) was recorded. The *p-*value for the cluster found in the observed data was calculated as the proportion of clusters selected from the permuted data sets that were larger than the observed cluster size. This corresponds to a test of the one-sided null-hypothesis that a positive serial dependence bias of the size observed in the participants’ behavior could have arisen by chance.

To investigate whether the strength of the serial dependence bias depended on the level of sensory uncertainty in cortex, we selected those orientation differences for which participant behavior exhibited a significant serial dependence bias (i.e., 1-57°), and divided these data (independently for each observer) into two bins. The first bin contained trials for which uncertainty (decoded from cortical activity) was higher on the current than previous trial, and the second bin consisted of trials for which decoded uncertainty was higher on the previous than current trial. Bias magnitude was calculated for an identical distribution of orientation differences in each of the two bins, and then averaged across observers. To assess significance, individual-observer biases were averaged across orientation differences, and a paired t-test was used to determine whether there was a significance difference between the two bins.

To determine whether the naturalistic and temporally misinformed observer models explained a statistically significant degree of variance in the participants’ group-average behavior, we used permutation tests. To assess the degree to which the observer model significantly captured serial dependence biases per se, we compared its observed goodness-of-fit (*R*^2^) to a null distribution of *R*^2^-values. This null distribution was generated from 10,000 random permutations of the observed data. Each permutation was computed by randomly shuffling the errors in the participants’ orientation reports with respect to the trial labels (i.e. the difference in stimulus orientation between consecutive trials). This simulates the null hypothesis that the behavioral errors arose from a distribution with constant mean, rather than one that depended on the orientation difference between trials (i.e. the behavioral bias does not depend on the difference in orientation between consecutive trials under the null hypothesis). For each permuted data set, a smoothed and group-averaged serial dependence curve was computed. This curve was then fit with the observer model, and the *R*^2^-value of this fit was recorded. The *p*-value of the observed *R*^2^ was computed as the proportion of permutations for which the *R*^2^-value was larger than the observed *R*^2^. This corresponds to a test of the null-hypothesis that the model provides a good fit of the serial dependence curve by chance.

A similar procedure was used to determine whether a naturalistic observer model accounting for trial-by-trial fluctuations in sensory uncertainty better captured the data than one that ignored these fluctuations and instead assumed one level of uncertainty in its decisions. Data were sorted into two bins, based on the trial-by-trial uncertainty levels estimated from the fMRI activity patterns (see also above). The first bin contained trials for which uncertainty was higher on the current than previous trial, and the second bin consisted of trials for which uncertainty was lower on the current than previous trial. The null distribution for this analysis was simulated by re-dividing trials into two equal-sized bins at random (rather than based on uncertainty), for each of 10,000 randomizations. For each permuted data set, a smoothed and group-averaged serial dependence curve was computed. The two models were then fitted to the permuted and the observed data, and their difference in *R*^2^ was recorded for each data set. The *p*-value for the improvement in fit due to modeling fluctuations in uncertainty was computed as the proportion of *R*^2^ differences obtained from the permuted data sets that were larger than the observed change in *R*^2^ value. This corresponds to a test of the null-hypothesis that the observed improvement in *R*^2^ for the uncertainty-based model arose by chance. Note that the permutation test automatically correct for increases in model complexity, as the comparison is between permuted and observed data, and any advantage due to added degrees of freedom applies to both sets of data.

### Code availability

Custom code written in MATLAB will be made available upon request.

## Results

### The ideal observer

How best to make sensory inferences in an environment that is largely stable across time? We will first examine ideal observer behavior, and later compare the predictions from this normative model to the behavior of human observers. We focus on a task in which the observer estimates from noisy sensory inputs the orientation of simple stimuli that are presented in sequence. Two basic principles underlie the computations by which the ideal observer performs this task. First, the ideal observer has knowledge about the temporal correlations between successive stimuli in the environment (**Fig. 2**). This enables the observer to predict the upcoming stimulus, based on perceptual estimates of previous stimuli. Second, the ideal observer realizes that sensory information is uncertain: because sensory inputs are noisy, each measurement is consistent with a range of possible perceptual interpretations. Utilizing knowledge about uncertainty enables the observer to make more accurate decisions, by prioritizing sensory information that is more certain.

These two normative principles give rise to an optimal decision-making algorithm, illustrated in **Fig. 3** (and discussed in more detail in the *Methods* section). At the start of the decision-making process, the observer has some sensory knowledge about the most recently viewed orientation. To capture the degree of uncertainty in this knowledge, the observer represents this information as a probability distribution over all possible stimulus orientations. The wider this distribution, the larger is the range of orientations consistent with the sensory input, and thus the greater is the degree of uncertainty that the observer has about the inputs. This probability distribution is combined with an internal model of the temporal orientation statistics of the environment. This enables the observer to generate a prediction about which orientation the stimulus is likely to have next. The prediction is also represented by a probability distribution. The observer subsequently takes another sensory measurement, which provides additional information about stimulus orientation. This sensory information is combined with the observer’s prediction, arriving at a final probability distribution that optimally reflects all the available information about the current stimulus from both past and present sensory inputs. The observer reports the mean of this distribution as their estimate of the current stimulus orientation.

**Figure 3:**
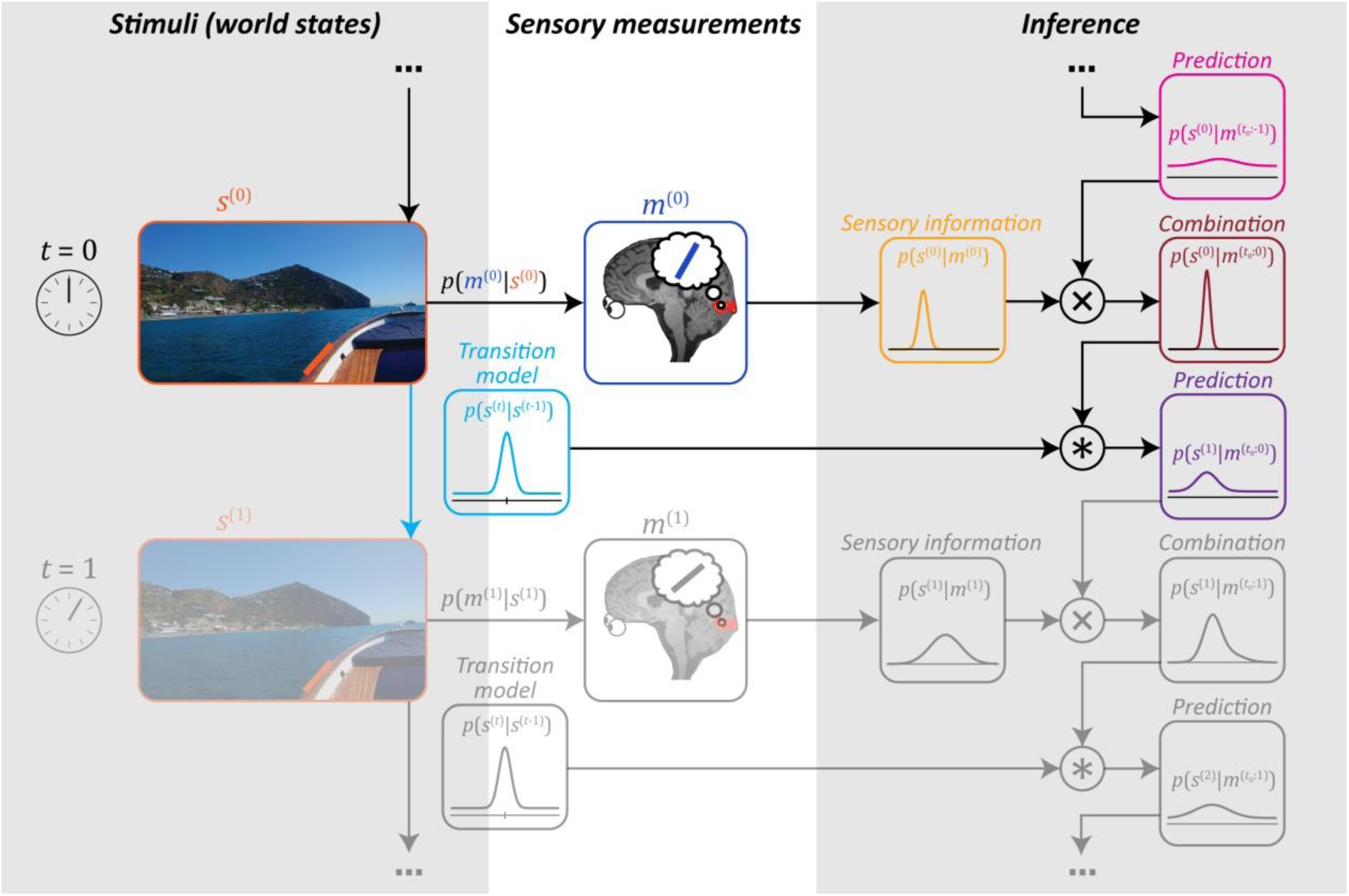
Ideal observer in the natural environment. The illustration highlights a single iteration in a continuous perceptual inference cycle. The left column illustrates the degree to which the environment remains stable over time, described by the transition model (light blue). For instance, edges in the environment (such as the one marked in orange) typically do not change much from one moment to the next. The ideal observer uses knowledge of these transition probabilities to infer the stimulus from its noisy sensory inputs. Specifically, at time *t* = 0, the observer takes a sensory measurement *m*^(0)^ of the stimulus *s*^(0)^ (an oriented edge). The measurement carries information about the stimulus, which is expressed by the probability distribution *p*(*s*^(0)^|*m*^(0)^) (yellow). This distribution is subsequently combined with a prediction (pink), which is based on previous sensory observations combined with knowledge of the environment’s temporal statistics. The prediction is expressed by a probability distribution 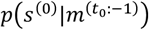, where *t*_0_ denotes the starting point of the inference process that is arbitrarily long ago. The prediction is integrated with incoming sensory information to arrive at a combined distribution 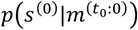 (red), which reflects all sources of knowledge available to the observer at *t* = 0. Finally, the observer uses this perceptual knowledge available at time 0 to generate a new prediction 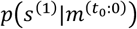 (purple) about the next stimulus at time *t* = 1, and the cycle continues.

To illustrate ideal observer behavior and the role of these various sources of knowledge, we simulated the observer’s responses to naturalistic sequences of oriented stimuli. Specifically, we used sequences of stimuli that followed naturalistic temporal statistics, measured using data obtained from real-world movies (see **Fig. 2**). We quantified the model observer’s performance as the average error in its orientation estimates, and compared this to the performance of three alternative model observers. Each of these observers was deficient in an important aspect of the ideal inference process. The first deficient observer ignored previous sensory inputs altogether, relying only on recent sensory information. The second deficient model observer ignored moment-to-moment changes in sensory uncertainty, while the third used an incorrect model of the temporal statistics of the environment by automatically integrating consecutive observations regardless of the change in orientation. We refer to these three as the ‘naïve’, ‘uncertainty-blind’ and ‘temporally misinformed’ observer, respectively. Comparing between the four model observers showed that none of the deficient observers matched the performance of the ideal (naturalistic) observer (**Fig. 4a**). This showcases the behavioral advantage of combining sensory estimates, as long as the integration process uses accurate knowledge of sensory uncertainty and environmental statistics.

**Figure 4:**
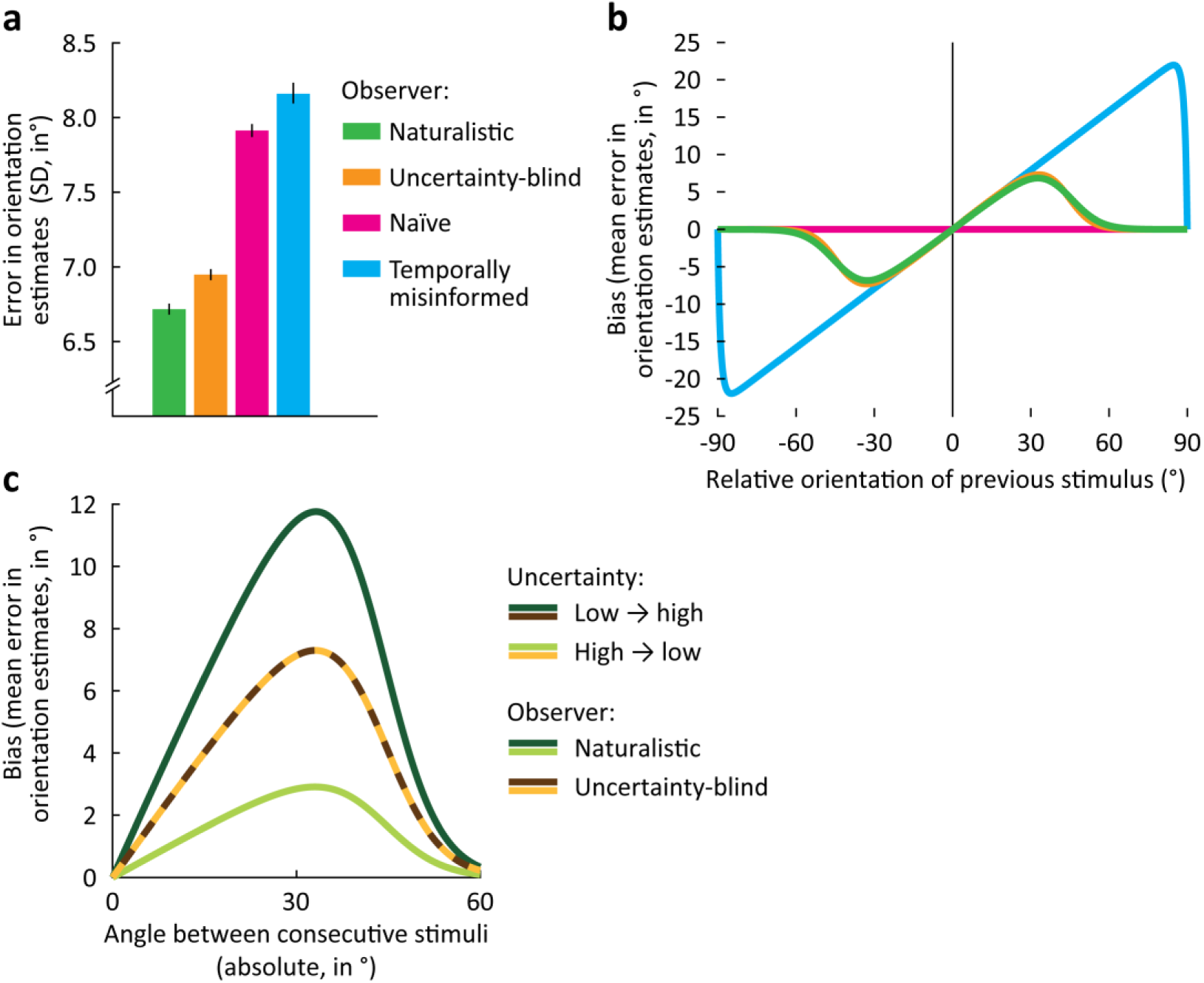
Simulated behavior of various observer models in an orientation estimation task. (**a**). Error (standard deviation) in orientation estimates for each of four model observers on sequences of orientation stimuli that were generated using naturalistic temporal statistics (i.e. correlations over time). The ideal or naturalistic observer utilizes knowledge of temporal stimulus statistics, while taking into account the uncertainty associated with successive stimulus estimates. The uncertainty-blind observer similarly combines previous and current sensory estimates, but fails to weight each by its associated uncertainty. The naïve observer bases estimates solely on sensory input from the current trial. The temporally misinformed observer takes into account uncertainty, but assumes an incorrect model of temporal statistics. The naturalistic observer outperforms the other three model observers that each ignore important aspects of the optimal inference process. Error bars depict bootstrapped 95% confidence intervals. (**b**) When presented with stimuli that are uncorrelated over time, the naturalistic observer incorrectly assumes temporal dependencies, and as a result, gives estimates that are biased towards previously seen stimuli (green). The same is true for the uncertainty-blind observer (orange), whose biases are nearly identical to those of the naturalistic observer (when averaged across uncertainty levels, as in this plot). This is in contrast to an observer model that assumes uncorrelated stimuli (the naïve observer); the estimates of this observer model exhibit no biases at all (pink). The temporally misinformed observer, on the other hand, does produce biased estimates, but combines current and previous estimates regardless of their difference in orientation (blue). See **Extended data figure 4-1** for a detailed analysis of model parameters and their effects on predicted behavior. For this and next panel, plotted is the predicted bias in the orientation estimates of the model observer as a function of the orientation of the previous relative to the current stimulus (acute-angle difference between previous and current orientation). For both axes, positive angles are in a clockwise direction. (**c**) For the uncertainty-aware naturalistic model observer, the bias is stronger when the observer is less uncertain about the previously seen stimulus than the current stimulus (dark green), than when the current sensory information was most reliable in comparison (light green). For the uncertainty-blind observer, the bias is identical across levels of uncertainty (brown and yellow).

Bringing the simulations closer to the actual conditions in which human observers were tested, we next examined the model observer’s behavior for sequences consisting of random orientation stimuli. Following previous work (Girshick et al., 2011; Fischer and Whitney, 2014), we assume that the naturalistic observer automatically applies knowledge about natural temporal statistics in their decisions, even when this internal world model provides no information about future inputs at all. While this strategy is certainly suboptimal in many experimental settings, it comes with substantial functional benefits in the generally stable, natural environment (**Fig. 4a**). The mismatch between actual experimental statistics and those assumed by the naturalistic observer created a substantial drop in behavioral accuracy (mean absolute error rose from 5.8° to 8.5°). Interestingly, we also observed a very distinctive pattern of behavioral errors that directly resulted from the observer’s sensory integration strategy. First, because the naturalistic observer operates under the assumption that consecutive inputs tend to arise from the same stimulus, behavioral orientation estimates were biased towards previously seen stimuli (**Fig. 4b**). Second, because the naturalistic observer uses knowledge about natural temporal dynamics, this bias peaks at relatively small stimulus differences (**Fig. 4b**). Third, because the naturalistic observer weights sensory inputs by their uncertainty, the temporal prediction has a stronger influence when current sensory information is comparatively unreliable. Consequently, the magnitude of the serial dependence effect is larger when a high-uncertainty trial follows a low-uncertainty trial (*low* → *high uncertainty*), compared to trials where uncertainty was greater on the previous trial (*high* → *low uncertainty*) (**Fig. 4c**). With these three predictions in hand, we now turn to human behavior to both test these predictions and, in doing so, probe the cortical representation of sensory uncertainty.

### Human observers

Do human observers integrate sensory evidence over time, while utilizing knowledge of natural temporal statistics and sensory uncertainty? To address this question, 18 participants performed an orientation estimation task while their brain activity was recorded with functional MRI. Orientation stimuli were presented in random sequence, and observers reported the orientation of each stimulus by rotating a bar presented at fixation. We compared human behavior against that of a model observer who assumes naturalistic statistical structure in the task. Specifically, we tested three predictions: (1) behavioral orientation estimates should be biased towards previously seen stimuli, (2) this bias should have a specific shape that reflects the temporal orientation statistics of the natural environment, peaking at small orientation differences, and (3) the magnitude of the bias should depend on the comparative reliability of current and previous sensory information.

To test for serial dependence in behavior, we compared the orientation reported on the current trial to that of the preceding trial. This comparison revealed that behavioral estimates were biased towards the orientation of the previous stimulus (cluster-based permutation test: *p* = 0.017; **Fig. 5a-b**), as predicted by the naturalistic observer model and observed in previous studies (e.g. Fischer and Whitney (2014); Fritsche et al. (2017)). To further quantify the effect, the naturalistic observer model was fit to participant behavior (see *Materials and Methods*). The model explained substantial and significant variance, suggesting that it captured relevant aspects of human behavior (*R*^*2*^ = 0.86, *p* = 0.02; parameter estimates: 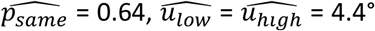, width (s.d.) and kurtosis of central peak in transition model: 16.9° and 2.6, respectively). These results were similar when serial dependence biases were computed with respect to the reported, rather than the presented orientation on the preceding trial, with a reliable bias over the same range of orientation differences (cluster-based permutation test: *p* = 0.002) that was well captured by the naturalistic observer model (*R*^2^ = 0.82, permutation test: *p* = 0.03). This suggests that human observers capitalize on the stability of the natural environment, by integrating current with previous sensory observations.

**Figure 5:**
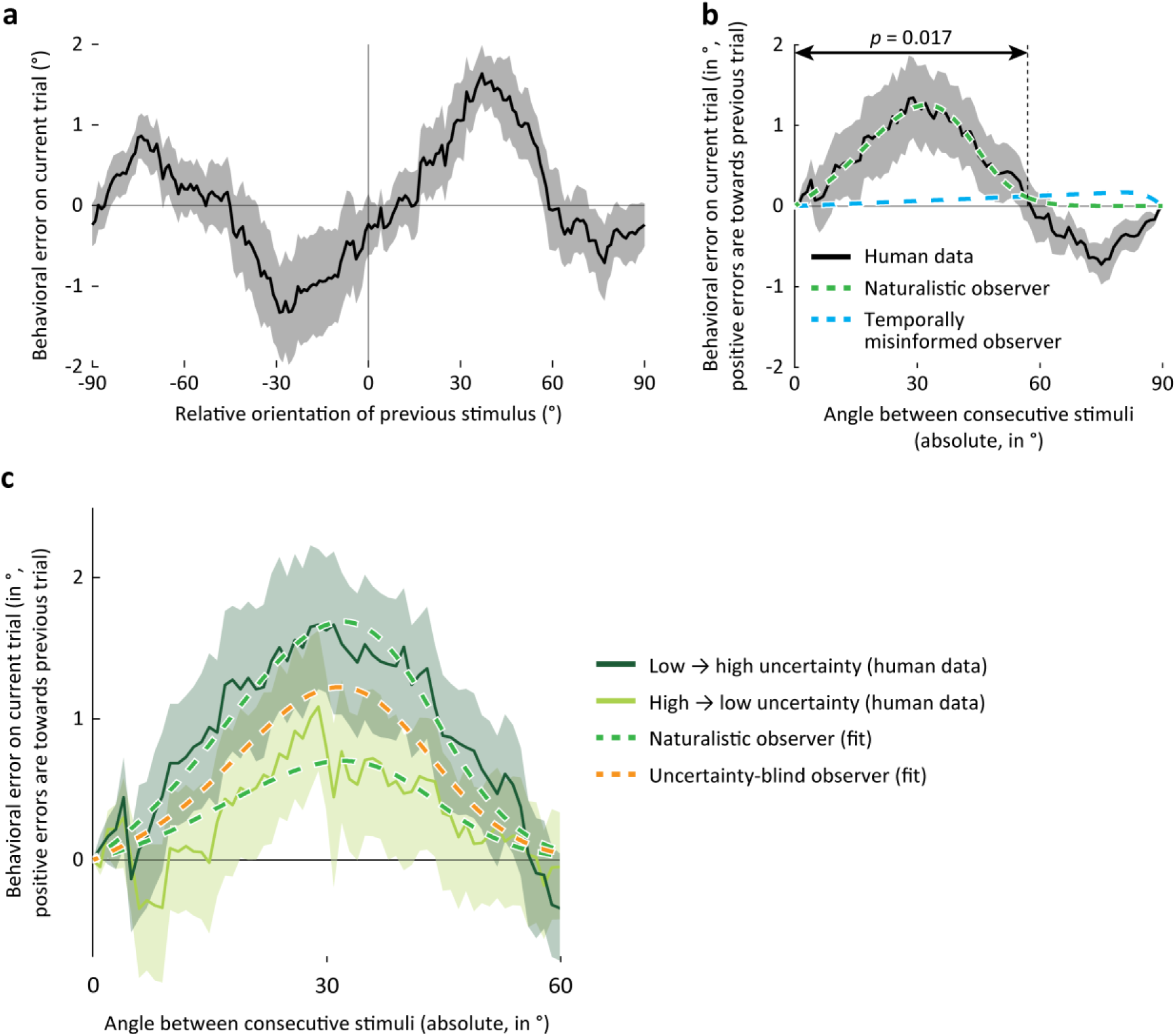
Human behavior matches the predictions of the naturalistic observer model. (**a**) Group-average behavioral errors, smoothed with a moving average filter, as a function of the relative orientation presented on the previous trial. Positive errors and orientation differences are in the clockwise direction. In this and subsequent panels, solid line and shaded region represent the mean +/-1 SEM across observers. (**b**) As in (**a**), but with behavioral errors realigned such that positive deviations are in the direction of the orientation presented on the previous trial, and plotted against the absolute, rather than the signed, difference in orientations. Arrows indicate the extent of orientation differences for which a cluster-based analysis revealed a significant bias in the direction of the previous trial (*p* = 0.017). A small negative serial dependence bias is also apparent for orientation angles larger than ± 60°, but did not reach statistical significance (*p* = 0.079, cluster-based permutation test). The green curve shows the optimal fit of the naturalistic ideal observer model to the behavioral data. This model explains significant variance in the data (*R*^*2*^ = 0.86, *p* = 0.02). The blue curve shows the optimal fit of an alternative observer model ignoring natural temporal statistics. The bias curve generated by this model is very different in shape from that observed in human behavior, and does not capture the human data well (*R*^2^ = 0.02, *p* = 0.41). (**c**) Similar to the naturalistic observer, the serial dependence bias in human behavior is stronger when the sensory representation is *more* uncertain on the current than previous trial (dark green), compared to when sensory information is *less* uncertain now than in the recent past (light green; *t*(17) = 2.96, *p* = 0.009). Dashed curves show the optimal fit of the naturalistic observer (green) and uncertainty-blind observer models (orange). The naturalistic observer model explained significantly more variance than a model ignoring fluctuations in uncertainty (*R*^*2*^ increase from 0.65 to 0.78, *p* = 0.01), suggesting that like the naturalistic observer model, human observers integrate current and previous sensory inputs in an uncertainty-weighted manner.

To test the degree to which knowledge of natural temporal statistics is necessary to capture the behavioral effect, we next compared human behavior against an observer model that operated under the naïve assumption that consecutive sensory observations always arise from a common source. Given this assumption, the optimal integration strategy is to simply average uncertainty-weighted sensory observations over time, with no regard to their orientation difference (as implemented by the temporally misinformed observer model). Note that human observers appear to compute such linear averages in many other sensory integration contexts (Ernst and Banks, 2002; Knill and Saunders, 2003; Alais and Burr, 2004). Interestingly, the temporally misinformed observer model failed to appropriately capture the shape of the serial dependence curve (*R*^*2*^ = 0.02, *p* = 0.41; **Fig. 5b**): while this model predicts a stronger bias the larger is the orientation difference between current and previous observations, serial dependence effects in participant behavior reached a maximum at relatively small orientation differences – a pattern that was well captured by the naturalistic observer model (see above). This suggests that human observers do not merely average sensory observations, but rather utilize an internal model of natural temporal statistics to determine how successive sensory inputs are best combined.

Finally, we examined whether human behavior is consistent with a Bayesian sensory integration strategy that takes into consideration sensory uncertainty. We reasoned that if uncertainty is represented in cortical stimulus representations and utilized in decisions, then this cortical representation of uncertainty should be predictive of trial-by-trial changes in the magnitude of the behavioral serial dependence bias. To characterize the degree of sensory uncertainty associated with cortical stimulus representations in areas V1-V3, we used a recently developed probabilistic decoding algorithm (van Bergen et al., 2015; van Bergen and Jehee, 2018) that calculates for each trial of cortical activity a posterior probability distribution over stimulus orientation. The width of this distribution can be taken as a metric of the amount of uncertainty contained in cortical activity. For each observer, trials were divided into two bins: one bin with trials for which the decoded cortical uncertainty was higher on the current compared to the previous trial (*low* → *high uncertainty*), and a second bin for which the width of the decoded probability distribution was narrower on the current compared to the previous trial (*high* → *low uncertainty*). As no external noise was added to the stimuli, binning was based solely on the degree of uncertainty in cortical activity patterns. We compared between behavioral biases for each of the two bins, predicting that their strength should change if observers utilized the degree of uncertainty associated with each of the two cortical stimulus representations.

Does the magnitude of the behavioral serial dependence bias depend on the reliability of the observer’s internal sensory evidence? Interestingly, behavioral orientation reports tended to be more biased towards the recent past when previous information in visual cortex was more reliable (*t*(17) = 2.96, *p* = 0.009; **Fig. 5c**), suggesting that human observers take into account the uncertainty in their cortical representations. Fitting the naturalistic observer model to the data, and comparing to the uncertainty-blind observer model, similarly showed that human behavior was better explained with a model that takes trial-by-trial fluctuations in uncertainty into account (significant increase in *R*^*2*^, from 0.65 to 0.78; permutation test on the difference in *R*^*2*^: *p* = 0.01; parameter estimates for the naturalistic observer: 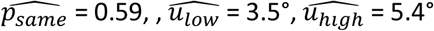, width (s.d.) and kurtosis of central peak in transition model: 17.3° and 2.6, respectively). The results were qualitatively similar when serial dependence biases were computed with respect to the reported, rather than presented, orientation on the preceding trial: as before, behavioral biases were reliably stronger when decoded uncertainty was higher on the current compared to the preceding trial (paired *t*-test: *t*(17) = 2.47, *p* = 0.024), and the naturalistic observer model explained more variance in behavior than the uncertainty-blind observer model (naturalistic observer: *R*^2^ = 0.76, uncertainty-blind observer: *R*^2^ = 0.63, permutation test on the difference in *R*^2^: *p* < 0.001). Together, these results suggest that human observers not only capitalize on the temporal continuity of the natural word by integrating sensory information across time, but also weight each piece of temporal evidence by its uncertainty. Moreover, it appears that the uncertainty that observers utilize in their perceptual decisions is directly linked to the cortical representation of the stimulus itself, providing critical evidence for probabilistic theories of neural coding (Zemel et al., 1998; Hoyer and Hyvärinen, 2003; Jazayeri and Movshon, 2006; Ma et al., 2006; Fiser et al., 2010).

## Discussion

How does the brain represent the reliability of its sensory evidence? Here, we tested an assumption central to Bayesian models of neural coding: that information is represented in cortical activity as a probability distribution, the width of which reflects the observer’s uncertainty. In testing these theories, we capitalized on a well-known behavioral bias called serial dependence, demonstrating that serial dependence in behavior is consistent with a statistical inference process that takes advantage of a temporally predictable natural environment. The modeling work resulted in quantitative predictions regarding sensory uncertainty that we tested with fMRI and a probabilistic decoding analysis. Our fMRI findings directly corroborate Bayesian theories by showing that the fidelity of a cortical stimulus representation, extracted as the width of a probability distribution, is linked to perceptual decisions. This suggests that sensory uncertainty is not only represented in visual activation patterns, but also read out by downstream areas to improve perceptual decision-making.

Earlier work on the cortical code for uncertainty (van Bergen et al., 2015) relied on a poorly understood behavioral bias with ill-defined links to uncertainty, leaving room for alternative explanations. This earlier work focused on a repulsive bias away from the cardinal axes; however, it is currently unclear how this cardinal repulsion bias might benefit the observer as an inference or decision strategy. In contrast, the serial dependence effect studied here better lends itself for an explanation based on normative principles, given the availability of real-world videos to characterize the temporal statistics of the natural environment. The here-discussed model implements a Bayesian recursive estimation strategy, whereby sensory estimates are continuously updated based on both knowledge of natural temporal orientation statistics and an uncertainty-weighted combination of current and previous observations to make the best possible decisions. The close match between model and human behavior suggests that serial dependence in perception (Fischer and Whitney, 2014) can be understood in relation to the features of such a recursive estimation process, and provides strong support for the hypothesis that the imprecision in a cortical stimulus representation reflects Bayesian uncertainty or probability.

Previous work on Bayesian inference has used external sources of noise, such as stimulus blur or contrast, to manipulate uncertainty. This is problematic because it could be that observers simply monitor such image properties as external cues to uncertainty. Compounding the issue, variations in physical stimulus properties typically affect cortical activity, making it impossible to discern whether cortical responses reflect a change in sensory uncertainty *per se* or rather one in stimulus features. For this reason, we held physical stimulus properties constant, and relied on internal fluctuations in activity to make perceptual information more or less reliable to the observer. We showed that uncertainty in cortical representations is directly linked to the uncertainty that human observers appear to use in their decisions, suggesting that downstream areas have access and utilize the reliability of early-level stimulus representations. Precisely how this information travels through the brain and modulates downstream decisional stages is an interesting question for future research.

Our results have important implications for theories of neural encoding. Specifically, while uncertainty may be represented by dedicated neural populations in e.g. the dopaminergic system (Fiorillo et al., 2003), our findings make it unlikely that specific visual cortical neurons represent summary statistics or are tuned to sensory uncertainty (Vilares and Körding, 2011). Rather, it appears that observers rely on the reliability of the cortical stimulus representation itself, suggesting that the same neural populations that represent the stimulus also carry information about its uncertainty. This direct link between the cortical representation of a stimulus and that of uncertainty supports encoding schemes such as probabilistic population codes (Ma et al., 2006), or sampling-based representations of probability (Hoyer and Hyvärinen, 2003; Fiser et al., 2010). It will be interesting for future neurophysiological studies that afford greater temporal resolution to dissociate between these and other alternatives for how probability distributions are encoded in neural activity.

Our findings suggest that human observers utilize knowledge of natural temporal statistics to determine whether or not previous sensory observations should be integrated, or rather segregated, in the current decision. Such an ideal serial integration process predicts biases towards recently seen stimuli when the difference between successive stimuli is relatively small, much like we observed in participant behavior. Interestingly, for larger changes in orientation (orientation angles larger than ± 60°), we additionally observed a tentative repulsive bias *away* from the previous stimulus, although this effect was only marginally significant. Such a repulsive effect at extreme stimulus differences has been reported by a few previous behavioral studies on serial dependence (Bliss et al., 2017; Fritsche et al., 2017), though not by others (Fischer and Whitney, 2014; St. John-Saaltink et al., 2016), and appears to reflect a different neural process that operates in parallel with the attractive bias considered here (Fischer & Whitney, 2014; Schwiedrzik et al., 2014; Fritsche et al., 2017; Kiyonaga et al., 2017). For example, whereas the attractive effect transfers across retinotopic locations, the repulsive effect is spatially specific, suggesting that it arises due to a relatively low-level process akin to sensory adaptation (Fischer & Whitney, 2014; Fritsche et al., 2017). Moreover, attractive and repulsive biases evolve in different directions during working memory maintenance (Bliss et al., 2017; Fritsche et al., 2017), and appear to map onto distinct cortical networks (Schwiedrzik et al., 2014). While the current study focuses on attractive effects in behavior, the model does not preclude any additional influences on serial decisions, which may, for instance, work towards suppressing redundant information or detecting change in the environment (Schwiedrzik et al., 2014). It will be interesting for future research to disentangle this interplay between positive and negative serial dependencies in perceptual decisions.

Our work is related to previous studies on predictions in perception. Several studies examined the behavioral and neural correlates of temporally constant perceptual priors (or expectations) in speed, direction of motion, or orientation perception (Weiss et al., 2002; Stocker and Simoncelli, 2006; Girshick et al., 2011; Kok et al., 2013; Vintch and Gardner, 2014). Our work differs in that we focus on the temporal predictability of sensory inputs from one moment to the next. Others have suggested that serial dependence effects in perception might reflect an advantageous sensory integration strategy employed by the brain to improve behavior (Cicchini et al., 2014, 2017, 2018; Fischer and Whitney, 2014). Very few of these studies, however, have cast this notion in an explicit normative framework. One notable exception (Cicchini et al., 2014) proposed a Kalman filter-like model that based its predictions on not only previous sensory inputs, but also their associated uncertainty. We extend this work by incorporating an explicit model of real-world temporal statistics in perceptual predictions, and utilize the framework to investigate how uncertainty is represented in visual cortex. Our findings indicate that human observers temporally combine sensory inputs in a statistically advantageous fashion by relying on the precision of internal stimulus knowledge. More fundamentally, our results advance understanding of how the nervous system represents uncertainty by showing that the fidelity of a cortical stimulus representation is directly linked to the uncertainty that observers appear to use in their decisions.

It is interesting to note that a model observer that simply averaged uncertainty-weighted sensory observations over time failed to capture human behavior. Instead, behavioral biases were well described by a model observer that combines sensory observations based on an internal model of the temporal statistics in the natural environment. Our work, in this sense, is similar in spirit to previous behavioral studies on motion tracking and (sensori)motor control, that have likewise found that human observers appear to employ an internal model of external world dynamics in motor decisions (Wolpert et al., 1995; Mehta and Schaal, 2002; Xivry et al., 2013; Kwon et al., 2015). Interestingly, although the description of the environment’s temporal dynamics used here was necessarily rather simplified, the model nonetheless captured several important aspects of human behavior. This suggests that the description, albeit it simple, still reflected some of the dominant features (e.g., general stability mixed with occasional unpredictability) of the full (and likely more complex) internal world model used by human observers, at least within the context of a simple orientation perception task.

Our findings are compatible with recent decisional accounts of serial dependence (e.g. Bliss et al. (2017); Fritsche et al. (2017), but see Cicchini et al. (2017); Fornaciai and Park (2018)) that propose that not perception itself, but rather later decisional stages bias behavior towards previously seen stimuli. Specifically, the ideal observer framework proposed here capitalizes on the stability of the visual world to make the best possible decisions, integrating sensory observations across time while weighting each piece of evidence by its uncertainty. It seems likely that this integration process is implemented in downstream areas involved in decision-making or working memory (Bliss et al., 2017; Kalm and Norris, 2018) that receive and retain various pieces of evidence over time, although our results do not rule out the possibility that visual representations are, in fact, combined at the earliest stages of sensory processing.

In conclusion, we found that serial dependence in behavior is consistent with a statistically advantageous sensory integration process that leverages the stability of the natural environment. Along with other findings in human orientation and speed perception (Weiss et al., 2002; Stocker and Simoncelli, 2006; Girshick et al., 2011) this suggests that certain behavioral biases, while seemingly maladaptive in experimental settings, are likely beneficial in the natural world. No less important, the observed link, both here and in our previous work (van Bergen et al., 2015), between cortical uncertainty and human behavior provides converging critical evidence for Bayesian theories of perception, suggesting that uncertainty is not only represented in cortical stimulus representations, but also utilized by downstream areas to refine perceptual decisions.

## Acknowledgements

This work was supported by ERC Starting Grant 677601 to J.J.

---

**Extended data figure 2-1:**
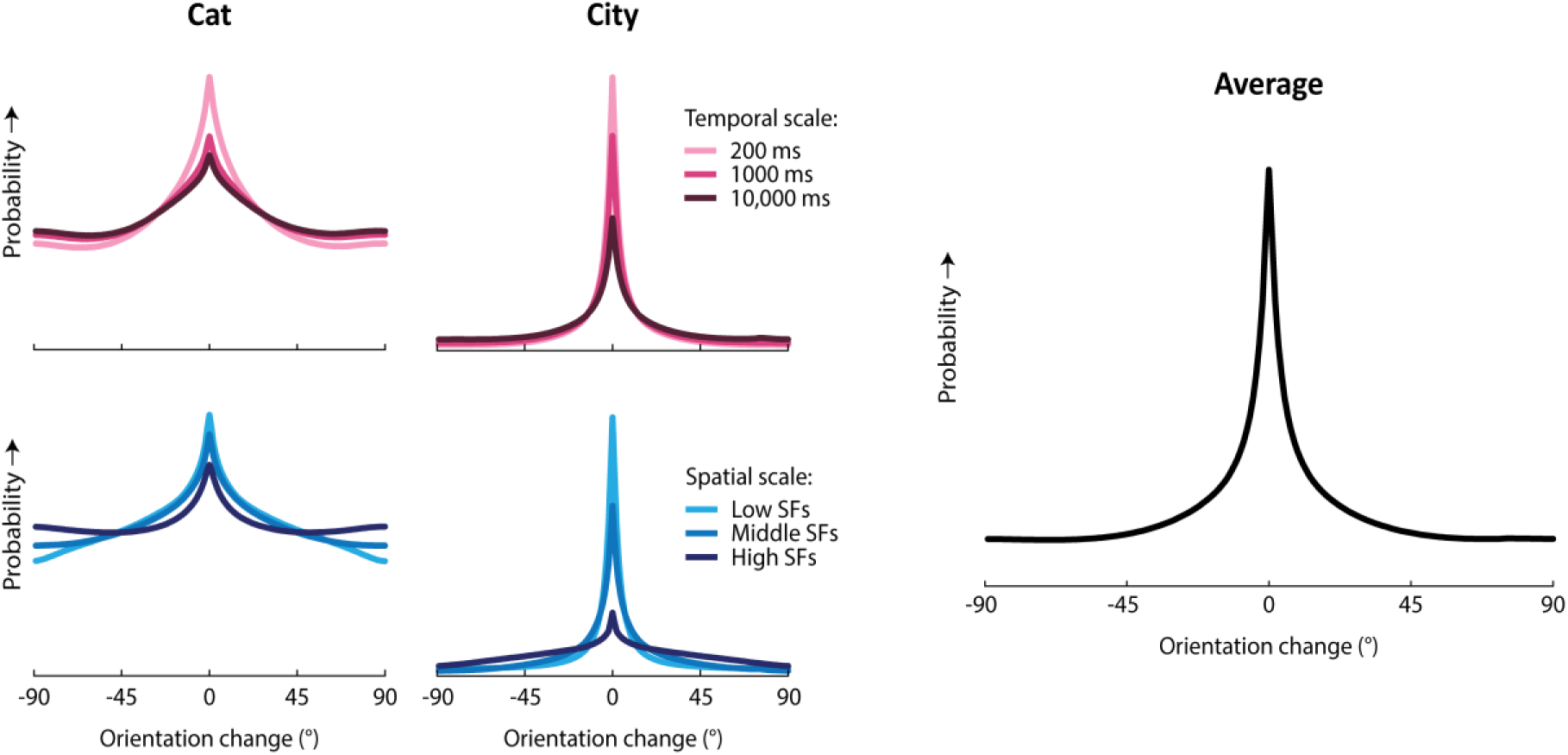
Temporal orientation statistics of natural videos.

Temporal orientation statistics were quantified using two databases of natural videos. The first database contained 17 grayscale videos shot through head-mounted cameras on cats roaming through various outdoor environments (Kayser et al., 2003; Betsch et al., 2004). Videos in this database had a spatial resolution of 320 × 240 pixels, a framerate of 25 Hz, and were recorded over temporal windows ranging from 38 to 200 seconds. The second database consisted of 18 color videos of outdoor scenes in a European city (Dorr et al., 2010). These videos were shot through mostly static cameras (with two videos containing minimal tracking motion), with spatial resolution 1280 × 720 pixels, framerate 30 Hz and duration between 19-20 seconds (Dorr et al., 2010). We will refer to these as the ‘cat’ and ‘city’ databases, respectively.

Videos were pre-processed by cropping each video to its central square (i.e. 240 × 240 and 720 × 720 pixels for *cat* and *city* videos, respectively), converting them to grayscale, and linearly normalizing the pixel intensities in each video frame to the [0, 1] range. The orientation content of each video was then characterized as follows. First, each video frame was filtered in the Fourier domain with a set of twelve orientation filters at three spatial scales. Specifically, the *k*-th orientation filter at the *l*-th spatial scale was defined as:

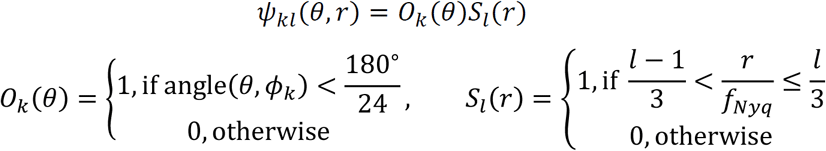

Here, *θ* and *r* are polar angle and eccentricity coordinates in Fourier space (corresponding to orientation and spatial frequency in the image domain). *ψ*_*kl*_ is the (*k, l*)-th Fourier filter, obtained by multiplying the *k*-th orientation filter *O*_*k*_(*θ*) with the *l*-th spatial frequency filter *S*_*l*_(*r*). Orientation and spatial frequency filters were boxcar functions. Orientation filters uniformly tiled orientation space, with the *k*-th filter centered on orientation 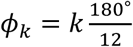. Spatial filters uniformly tiled the spatial frequency range between 0 and the Nyquist frequency *f*_*Nyq*_. Each double-boxcar Fourier filter was smoothed with a Gaussian kernel (standard deviation: 20 pixels) to prevent ringing artefacts in the filtered image.

This procedure separated the image into 36 Fourier components, each of which was then transformed back into the image domain. For each spatial scale *l*, image pixel *j* and time point *t*, this yields a vector 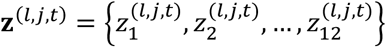, of twelve values describing the local energy in each orientation band. These orientation intensities were averaged in circular space to obtain a complex vector pointing towards their average orientation:

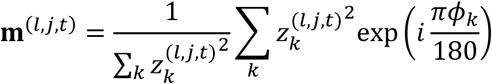

where *i* is the imaginary unit. The angle of this vector served as a measurement of the average orientation *μ*^(*l,j,t*)^ at spatial scale *l*, location *j* and time *t*, while its length *λ*^(*l,j,t*)^ quantified how tightly the orientation content was concentrated around this mean (such that higher values indicated a more pronounced orientation signal). Thus, for each video frame and spatial scale, this analysis yields a map that describes for each pixel its average orientation and orientation strength.

After characterizing the orientation content of each video frame, we measured its evolution over time, separately for each spatial scale. First, all pairs of video frames {*t*_1_, *t*_2_} were selected that were separated by a given time interval. To exclude noise, we then selected, for each pair of frames, all pixels with strong orientation content across both time points (frames). This was defined as those pixels for which the product 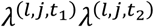 was in the top 50% of that particular pair of frames. For each of these pixels, we recorded their orientations 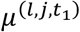 and 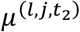 at the lagging and leading frames, respectively, and added these values to a histogram, characterizing the joint probability distribution of current and previous orientations for a given image location and spatial scale. One such histogram was obtained for every video, and a mean histogram was computed across all videos in each database. Finally, this two-dimensional joint distribution 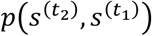 of current and preceding orientations was converted to a one-dimensional transition kernel. First, we conditioned the distribution on the preceding orientation, by computing 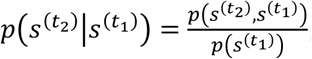, where 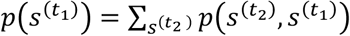. The joint distribution describes the overall probability that one orientation is followed by another, while the conditional distribution expresses the probability of the second orientation given a certain value of the first. We then collapsed this conditional distribution onto a single dimension, by recasting it in terms of the difference in orientation between 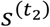 and 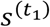. This results in a transition distribution, summarizing the probability that an orientation will change by a certain amount in a given timespan, regardless of the initial orientation. Transition distributions were estimated separately for each spatial scale, and for a range of time intervals (200, 1000, and 10,000 ms).

The figure summarizes the results of this analysis. It shows the orientation transition distribution measured in the two databases for different temporal intervals (averaged across spatial scales) and spatial scales (averaged across temporal intervals), as well as the overall mean across all spatial and temporal scales and both databases. Notice that the same overall shape appears regardless of temporal window, spatial scale or database: a central peak, comprising orientation changes of limited range, combined with a uniform component reflecting random changes in orientation. The specific parameters of this shape vary somewhat between videos and scales. The distribution tends to be wider, with a stronger uniform component, for smaller compared to larger spatial scales, and also for longer compared to shorter temporal intervals. This makes sense, as larger changes are likely to occur over longer timespans, and orientation content for high spatial frequencies is likely to change more quickly. On average, orientation varies also more quickly and randomly over space and time in the *cat* database, which may be explained by the abundant camera motion in these videos, compared to the mostly static viewpoints in the *city* database. Across databases and scales, the excess kurtosis of the distributions varied from -0.3 (i.e. roughly Gaussian) to 5.8, the circular standard deviation of the central peak ranged from 12.4-33.5°, and the contribution of the uniform component varied between 24-94% (comparable to *p*_*same*_ = 0.06-0.76).

Following these natural temporal statistics, the orientation transition kernel in the naturalistic observer model was similarly described as a mixture between a central peak and uniform component. The parameter values of the mixture distribution (i.e., the width of the peak, its kurtosis, and the contribution of the uniform component) were fit to the behavioral data, as natural transition distributions were found to vary in these dimensions across analysis settings (see above) and it is a-priori unclear which specific values would best describe the visual environment of the fMRI-experiment.

**Extended data figure 4-1:**
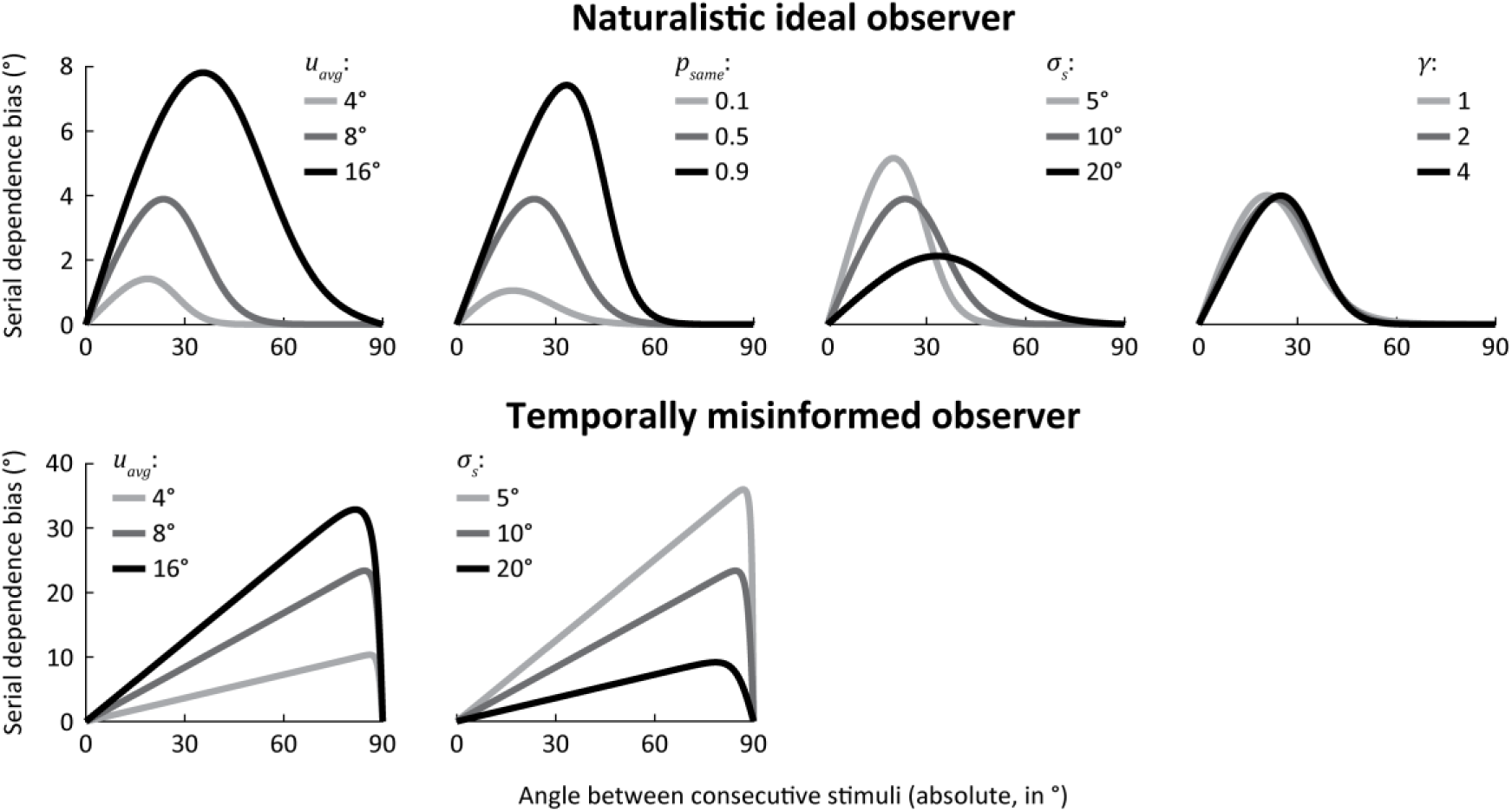
Behavioral repertoire of model observers.

To illustrate the range of serial dependence curves consistent with each of the different model observers, we simulated model behavior using various values of model parameters. The magnitude of the serial dependence bias on the current trial is plotted against the absolute difference in orientation between current and previous stimuli. By definition, the serial dependence bias of the naïve observer is always zero, regardless of orientation angle, and is not shown here.

Plots show the across-trial average of the serial dependence bias curve. Trial-by-trial fluctuations in uncertainty change the magnitude, but not the shape, of the bias curve, as can be seen in **Fig. 4c**. When averaged across uncertainty levels, the bias curve of the naturalistic observer is practically identical to that of the uncertainty-blind observer (see **Fig. 4b**), which is why the latter is not shown here.

Parameter values were manipulated around default settings of *u*_*avg*_ = 8°, *p*_*same*_ = 0.5, *σ*_*s*_ = 10° and *γ* = 2. In each plot, the dark gray curve corresponds to these default parameters, and is identical between panels (for the same observer). Parameters *p*_*same*_ and *γ* are not free parameters in the temporally misinformed observer model, and were not manipulated there. Note that for the temporally misinformed observer, the serial dependence bias always equals 0 when orientations differ by exactly 90°, whereas the naturalistic observer’s bias can reach 0 much earlier, depending on parameter values. This is because the temporally misinformed observer always averages together previous and current sensory observations, regardless of their orientation difference. Consequently, the serial bias of this observer is only 0 when the circular average of these orientations is 0 (i.e. when orientations are orthogonal).

The different values of *γ* corresponded to excess kurtoses of 3 (Laplacian), 0 (Gaussian) and -0.8 (sub-Gaussian), for *γ* = 1, 2, and 4 respectively. For the default setting of *γ* = 2, *σ*_*s*_ corresponds to the standard deviation of the central peak in the transition model. When *γ* is set to different values, however, this changes both the kurtosis and standard deviation of the central peak. To illustrate the specific effect of changing kurtosis, *σ*_*s*_ was therefore adjusted along with *γ* to keep the standard deviation fixed at an approximately constant value of 10°. As evident from the resulting plots, changing the kurtosis of the central peak does not strongly affect the shape of the serial dependence curve. This is because the transition distribution is convolved with a (roughly) Gaussian distribution (which reflects the observer’s knowledge about the previous stimulus). The result of this convolution is the observer’s prediction. For a reasonable range of kurtosis values of the transition distribution, this prediction always tends towards a Gaussian shape.

## References

Alais D, Burr DC (2004) The ventriloquist effect results from near-optimal bimodal integration. Curr Biol 14:257–262.

Anastasio TJ, Patton PE, Belkacem-Boussaid K (2000) Using Bayes’ rule to model multisensory enhancement in the superior colliculus. Neural Comput 12:1165–1187.

Appelle S (1972) Perception and discrimination as a function of stimulus orientation: the “oblique effect” in man and animals. Psychol Bull 78:266–278.

Battaglia PW, Jacobs RA, Aslin RN (2003) Bayesian integration of visual and auditory signals for spatial localization. J Opt Soc Am A 20:1391.

Betsch BY, Einhäuser W, Körding KP, König P (2004) The world from a cat’s perspective - Statistics of natural videos. Biol Cybern 90:41–50.

Bliss DP, Sun JJ, D’Esposito M (2017) Serial dependence is absent at the time of perception but increases in visual working memory. Sci Rep 7:14739.

Brainard DH (1997) The Psychophysics Toolbox. Spat Vis 10:433–436.

Brouwer GJ, Heeger DJ (2011) Cross-orientation suppression in human visual cortex. J Neurophysiol 106:2108–2119.

Chopin A, Mamassian P (2012) Predictive Properties of Visual Adaptation. Curr Biol 22:622–626.

Cicchini GM, Anobile G, Burr DC (2014) Compressive mapping of number to space reflects dynamic encoding mechanisms, not static logarithmic transform. Proc Natl Acad Sci 111:7867–7872.

Cicchini GM, Mikellidou K, Burr D (2017) Serial dependencies act directly on perception. J Vis 17:1–9.

Cicchini GM, Mikellidou K, Burr DC (2018) The functional role of serial dependence. Proc R Soc B Biol Sci 285:20181722.

DeYoe EA, Carman GJ, Bandettini P, Glickman S, Wieser J, Cox R, Miller D, Neitz J (1996) Mapping striate and extrastriate visual areas in human cerebral cortex. Proc Natl Acad Sci 93:2382–2386.

Dong D, Atick J (1995) Statistics of natural time-varying images. Netw Comput Neural Syst 6:345–358.

Dorr M, Martinetz T, Gegenfurtner KR, Barth E (2010) Variability of eye movements when viewing dynamic natural scenes. J Vis 10:28–28.

Engel SA, Glover GH, Wandell BA (1997) Retinotopic organization in human visual cortex and the spatial precision of functional MRI. Cereb Cortex 7:181–192.

Ernst MO, Banks MS (2002) Humans integrate visual and haptic information in a statistically optimal fashion. Nature 415:429–433.

Fiorillo CD, Tobler PN, Schultz W (2003) Discrete coding of reward probability and uncertainty by dopamine neurons. Science 299:1898–1902.

Fischer J, Whitney D (2014) Serial dependence in visual perception. Nat Neurosci 17:738–743.

Fischl B, Sereno MI, Dale AM (1999) Cortical surface-based analysis. II: Inflation, flattening, and a surface-based coordinate system. Neuroimage 9:195–207.

Fiser J, Berkes P, Orbán G, Lengyel M (2010) Statistically optimal perception and learning: from behavior to neural representations. Trends Cogn Sci 14:119–130.

Fornaciai M, Park J (2018) Attractive Serial Dependence in the Absence of an Explicit Task. Psychol Sci 29:437–446.

Fritsche M, Mostert P, de Lange FP (2017) Opposite Effects of Recent History on Perception and Decision. Curr Biol 27:590–595.

Furmanski CS, Engel SA (2000) An oblique effect in human primary visual cortex. Nat Neurosci 3:535–536.

Gibson JJ, Radner M (1937) Adaptation, after-effect and contrast in the perception of tilted lines. I. Quantitative studies. J Exp Psychol 20:453–467.

Girshick AR, Landy MS, Simoncelli EP (2011) Cardinal rules: visual orientation perception reflects knowledge of environmental statistics. Nat Neurosci 14:926–932.

Hoyer PO, Hyvärinen A (2003) Interpreting neural response variability as monte carlo sampling of the posterior. Adv Neural Inf Process Syst :293–300.

Jacobs RA, Fine I (1999) Experience-dependent integration of texture and motion cues to depth. Vision Res 39:4062–4075.

Jazayeri M, Movshon JA (2006) Optimal representation of sensory information by neural populations. Nat Neurosci 9:690–696.

Jenkinson M, Bannister P, Brady M, Smith S (2002) Improved Optimization for the Robust and Accurate Linear Registration and Motion Correction of Brain Images. Neuroimage 17:825–841.

Kalm K, Norris D (2018) Visual recency bias is explained by a mixture model of internal representations. J Vis 18:1.

Kayser C, Einhäuser W, König P (2003) Temporal correlations of orientations in natural scenes. Neurocomputing 52–54:117–123.

Kiyonaga A, Scimeca JM, Bliss DP, Whitney D (2017) Serial Dependence across Perception, Attention, and Memory. Trends Cogn Sci 21:493–497.

Knill DC, Saunders JA (2003) Do humans optimally integrate stereo and texture information for judgments of surface slant? Vision Res 43:2539–2558.

Kok P, Brouwer GJ, van Gerven MAJ, de Lange FP (2013) Prior expectations bias sensory representations in visual cortex. J Neurosci 33:16275–16284.

Kwon O-S, Tadin D, Knill DC (2015) Unifying account of visual motion and position perception. Proc Natl Acad Sci 112:8142–8147.

Liberman A, Fischer J, Whitney D (2014) Serial dependence in the perception of faces. Curr Biol 24:2569–2574.

Ma WJ, Beck JM, Latham PE, Pouget A (2006) Bayesian inference with probabilistic population codes. Nat Neurosci 9:1432–1438.

Maris E, Oostenveld R (2007) Nonparametric statistical testing of EEG- and MEG-data. J Neurosci Methods 164:177–190.

Mehta B, Schaal S (2002) Forward models in visuomotor control. J Neurophysiol 88:942–953.

Pelli DG (1997) The VideoToolbox software for visual psychophysics: transforming numbers into movies. Spat Vis 10:437–442.

Schwiedrzik CM, Ruff CC, Lazar A, Leitner FC, Singer W, Melloni L (2014) Untangling perceptual memory: Hysteresis and adaptation map into separate cortical networks. Cereb Cortex 24:1152–1164.

Sereno MI, Dale AM, Reppas JB, Kwong KK, Belliveau JW, Brady TJ, Rosen BR, Tootell RB (1995) Borders of multiple visual areas in humans revealed by functional magnetic resonance imaging. Science 268:889–893.

St. John-Saaltink E, Kok P, Lau HC, de Lange FP (2016) Serial Dependence in Perceptual Decisions Is Reflected in Activity Patterns in Primary Visual Cortex. J Neurosci 36:6186–6192.

Stocker AA, Simoncelli EP (2006) Noise characteristics and prior expectations in human visual speed perception. Nat Neurosci 9:578–585.

van Bergen RS, Jehee JFM (2018) Modeling correlated noise is necessary to decode uncertainty. Neuroimage 180:78–87.

van Bergen RS, Ma WJ, Pratte MS, Jehee JFM (2015) Sensory uncertainty decoded from visual cortex predicts behavior. Nat Neurosci 18:1728–1730.

Vilares I, Körding KP (2011) Bayesian models: the structure of the world, uncertainty, behavior, and the brain. Ann N Y Acad Sci 1224:22–39.

Vintch B, Gardner JL (2014) Cortical Correlates of Human Motion Perception Biases. J Neurosci 34:2592–2604.

Weiss Y, Simoncelli EP, Adelson EH (2002) Motion illusions as optimal percepts. Nat Neurosci 5:598–604.

Wolpert DM, Ghahramani Z, Jordan MI (1995) An Internal Model for Sensorimotor Integration. 269:1880–1882.

Xivry JO De, Blohm G, Lefe P (2013) Kalman Filtering Naturally Accounts for Visually Guided and Predictive Smooth Pursuit Dynamics. 33:17301–17313.

Zemel RS, Dayan P, Pouget A (1998) Probabilistic interpretation of population codes. Neural Comput 10:403–430.

Zhang W, Luck SJ (2008) Discrete fixed-resolution representations in visual working memory. Nature 453:233–235.

